# Priming of Myelin-Specific T Cells in the Absence of Dendritic Cells Results in Accelerated Development of Experimental Autoimmune Encephalomyelitis

**DOI:** 10.1101/2020.12.28.424609

**Authors:** Thaiphi Luu, Julie F. Cheung, Hanspeter Waldner

## Abstract

Experimental autoimmune encephalomyelitis (EAE), the animal model of multiple sclerosis (MS), is predominantly mediated by pro-inflammatory CD4^+^ T cell responses to CNS antigens, including myelin proteolipid protein (PLP). Dendritic cells (DCs) are considered critical for inducing T cell responses against infectious agents, but the importance of DCs in priming self-reactive CD4+ T cells in autoimmune disease such as MS has been unclear.

To determine the requirement of DCs in PLP-specific CD4^+^ T cell responses and EAE, we genetically deleted CD11c^+^ DCs in PLP T cell receptor (TCR) transgenic SJL mice constitutively. DC deficiency did not impair the development, selection or the pathogenic function of PLP-specific CD4^+^ T cells in these mice, and resulted in accelerated spontaneous EAE compared to DC sufficient controls. In addition, using a genetic approach to ablate DCs conditionally in SJL mice, we show that CD11c^+^ DCs were dispensable for presenting exogenous or endogenous myelin antigen to PLP-specific T cells and for promoting pro-inflammatory T cell responses and severe EAE. Our findings demonstrate that constitutive or conditional ablation of CD11c^+^ DCs diminished self-tolerance to PLP autoantigen. They further show that in the absence of DCs, non-DCs can efficiently present CNS myelin antigens such as PLP to self-reactive T cells, resulting in accelerated onset of spontaneous or induced EAE.

## Introduction

Experimental autoimmune encephalomyelitis (EAE) is a well-established animal model for multiple sclerosis (MS), a demyelinating autoimmune disease of the CNS [1–3]. Pro-inflammatory CD4^+^ T cells that are specific for CNS myelin antigens, including PLP, are the major mediators of EAE. The disease is actively induced in susceptible mice by immunization with myelin peptides in the presence of potent adjuvants such as CFA [4, 5]. Alternatively, transfer of myelin-specific CD4^+^ T cells into naive mice is an established method to induce passive EAE [6, 7].

Dendritic cells (DCs) represent the most potent antigen presenting cells (APCs) and are comprised of two major subsets, conventional DCs (cDCs) and plasmacytoid DCs (pDCs) [8]. Under steady-state conditions, DCs show an immature phenotype, which is characterized by low surface expression of MHC II and co-stimulatory molecules, including CD80, CD86 and CD40. Immature DCs induce peripheral tolerance by various mechanisms, including the expression of T cell regulatory factors such as indoleamine 2,3-dioxygenase (IDO) and IL-10, or by induction of regulatory T cells (Tregs) [9–11]. Following infection or inflammatory signals, including pro-inflammatory cytokines or microbial products, DCs up-regulate their expression of MHC II and co-stimulatory molecules and differentiate into mature DCs. Activated/mature DCs, which secrete inflammatory cytokines are proficient in priming and differentiating naive CD4^+^ T cells [12, 13].

The specific APCs that promote myelin-specific T cell responses and their precise role in MS and EAE pathogenesis have remained unclear [14–16]. We previously demonstrated that microbial activation of APCs, including DCs, broke CD4^+^ T cell tolerance to myelin PLP and triggered EAE in PLP TCR transgenic mice on a non-autoimmune background [17]. Our study and those by other laboratories supported a role of DCs in myelin-specific T cell activation and EAE induction [18–20]. Conversely, a number of other studies provided evidence that DCs can play a protective or ameliorative role in EAE [21–23]. More recently, Isaksson et al. challenged the importance of DCs in EAE pathogenesis by demonstrating that conditional ablation of cDCs had no significant effect on priming or differentiation of MOG-specific Th responses and mildly reduced EAE severity in C57BL/6 mice [24]. However, these results were in disagreement with the findings of a similar study by Yogef et al., which showed that DC ablation led to stronger inflammatory T cell responses and enhanced EAE severity [25]. Importantly, however, no study to date has addressed the role of DCs in the setting of spontaneous EAE, which develops in the absence of myelin antigen immunization.

Here, we generated two novel transgenic mouse lines on the autoimmune prone SJL strain, CD11c-DTR and PLP TCR Tg. CD11c-DTA mice, to ablate CD11c^+^ DCs conditionally and constitutively, respectively. For the first time, the latter mice allowed determining the requirement of DCs in the development of spontaneous EAE. In addition, CD11c-DTR mice enabled us to assess selectively the role of DCs in active and passive EAE in SJL mice. We demonstrate that constitutive or conditional depletion of DCs did not impair, but instead accelerated the induction of spontaneous and passive EAE, respectively. Furthermore, thymic development and selection of PLP-specific CD4^+^ T cells was not affected significantly by the lack of DCs in PLP TCR Tg. CD11c-DTA mice. Our findings indicate that DCs are not required in the pathogenesis of spontaneous or induced EAE, suggesting that non-DCs efficiently promote PLP-specific CD4^+^ T cell responses in the absence of DCs.

## Material and methods

### Mice

CD11c-Cre C57BL/6 mice (B6.Cg-Tg(Itgax-cre)1-1Reiz/J) [26], Rosa26-eGFP-DTA C57BL/6 mice (*Gt(ROSA)26Sor^tm1(DTA)Jpmb^*/J) [27], CD11c-DTR C57BL/6 mice (B6.FVB-*1700016L21Rik^Tg(Itgax-DTR/EGFP)57Lan^*/J) [12] were purchased from Jackson Laboratory and fully backcrossed onto SJL/J mice for ten generations at the Penn State College of Medicine animal facility.

SJL/J mice transgenic for the PLP T cell receptor (TCR) 5B6 specific for myelin PLP139-151, and recombination-activating gene 2 deficient (*Rag2*−/−) SJL mice had been generated previously and maintained at the Penn State College of Medicine animal facility [28]. PLP TCR transgenic SJL mice were bred with fully backcrossed Rosa26-eGFP-DTA or CD11c-Cre SJL mice. PLP TCR transgenic offspring that were transgenic for Rosa26-eGFP-DTA or CD11c-Cre served either as DC sufficient controls or were intercrossed with CD11c-Cre or Rosa26-eGFP-DTA SJL mice, respectively to generate DC deficient PLP TCR transgenic mice (called PLP TCR Tg. CD11c-DTA hereafter).

Fully backcrossed CD11c-DTR SJL mice were bred with *Rag2*−/− SJL mice. CD11c-DTR transgenic offspring were subsequently intercrossed with *Rag2*−/− SJL mice to generate CD11c-DTR *Rag2−/−* mice. Homozygosity for the *Rag2−/−* genotype was determined by PCR.

All mice were housed at the Penn State College of Medicine specific pathogen-free (SPF) animal facility in accordance with the guidelines of the Penn State Institutional Animal Care and Use Committee.

### Flow cytometry and intracellular cytokine staining

Single-cell suspensions of lymphoid tissues were prepared, and red blood cells were lysed by hypotonic shock. 1 × 10^6^ cells were stained with monoclonal antibodies (20 min, 4°C). The following monoclonal antibodies (BD Biosciences or eBioscience) were used for surface staining: anti-CD4 (RM4-5), anti-CD8α (53-6.7), anti-TCR Vβ6 (RR4-7), anti-CD45R/B220 (RA3-6B2), anti-CD11b (M1/70), anti-CD11c (HL3), anti-I-A^q^ (KH116, cross-reacts with I-A^s^), anti-CD69 (H1.2F3), anti-CD62L (MEL14), and anti-CD44 (IM7).

Absolute numbers of cell populations were calculated by multiplying the total cell count with the frequency of the respective population as determined by flow cytometry. CellTrace Violet staining was performed according to the manufacturer’s protocol (Invitrogen). For intracellular cytokine staining, axillary and brachial lymph node cells were cultured overnight in 24-well plates in culture medium (DMEM supplemented with 10% (v/v) FCS, 100 units/ml penicillin, 100 units/l streptomycin, 50 μM 2-mercaptoethanol, 10 mM HEPES, 1 mM sodium pyruvate, and 1 × MEM non-essential amino acids) in the presence of 25 μg/ml PLP139-151 (HSLGKWLGHPDKF, Abbiotec). The next morning, the cells were stimulated for 4 hrs with phorbol myristate acetate (PMA) (5 ng/ml) and ionomycin (500 ng/ml) in the presence of Brefeldin A (5 μg/ml) (all from Enzo Life Sciences). BD Cytofix/Cytoperm and Perm/Wash buffers were used for intracellular staining with the following monoclonal antibodies: anti-IFN-γ (XMG1.2), anti-IL-17 (TC11-18H10), IL-4 (11B11) or IL-10 (JESS-16E3) according to the manufacturer’s instructions (BD Bioscience or eBioscience). At least 3 × 10^4^ live cells, based on characteristic forward- and side-scatter profile gating, were acquired with a LSR II cytometer (BD). Data were analyzed with FlowJo software (BD).

### Depletion of dendritic cells in vivo

For conditional DC depletion, CD11c-DTR mice, CD11c-DTR BM or CD11c-DTR *Rag2−/−* BM chimeric mice were injected (i.p.) with 4-5 ng/g body weight diphtheria toxin (DT, Sigma-Aldrich) every other day for the indicated time. To assess the efficiency of DT induced DC ablation, single cell suspensions of lymphoid organs were prepared and treated (2 mg/ml, 30 min) with Collagenase D (Sigma-Aldrich) one day following the final injection of DT. Indicated lymphoid organs from PLP TCR Tg CD11c-DTA mice were processed similarly to determine the efficiency of constitutive DC ablation.

Single lymphoid cell suspensions were stained with monoclonal antibodies for indicated DC surface markers and analyzed by flow cytometry.

### Labeling and adoptive transfer of PLP-specific T cells

CD4^+^ T cells were enriched (> 95% purity) from spleen or axillary and brachial lymph nodes of naive PLP TCR transgenic SJL mice using magnetic beads according to the manufacturer’s instructions (Miltenyi Biotec). The cells were labeled with 3 μM CellTrace Violet (Invitrogen) according to the manufacturer’s instructions and injected (8 × 10^6^ cells/mouse; i.v.) into DT or PBS treated CD11c-DTR SJL mice. Immediately after T cell transfer, recipient mice were immunized (s.c.) with PLP139-151 emulsified in CFA (Difco) under each forelimb (12.5 μg) and into one hind footpad (3 μg). Four days later, PLP TCR transgenic (CD4^+^ TCR Vβ6^+^) T cells from draining lymph nodes of injected mice were analyzed for the dilution of CellTrace violet by flow cytometry.

### Generation of bone marrow chimeras

Bone marrow (BM) cells were harvested from femur and tibiae of 6–9 week-old CD11c-DTR or CD11c-DTR *Rag2−/−* SJL mice and depleted of T cells using anti-CD3 and anti-CD8 magnetic beads according to the manufacturer’s instructions (Miltenyi Biotec). Wild type or *Rag2−/−* SJL mice were gamma-irradiated with 950 rads from a ^60^Co source. The next day, irradiated mice were injected with T cell-depleted bone marrow cells (5-8 × 10^6^ cells/mouse) pooled from two to three donor mice. The recipient mice were maintained on water containing 1.5% (v/v) sulfamethoxazole/trimethoprim (200 mg/40 mg per 5 ml; Halo Pharmaceutical) for two weeks. They were analyzed for successful engraftment six to eight weeks after reconstitution before being used for experiments.

### Induction and assessment of EAE

To assess encephalitogenic function of PLP-specific T cells purified CD4^+^ T cells from naïve PLP TCR Tg. CD11c-DTA SJL mice or PLP TCR Tg. DTA control mice were transferred (1.2 × 10^6^ cells/mouse, i.v.) into *Rag2 −/−* SJL mice. To induce passive EAE, purified CD4^+^ T cells (3 × 10^6^ cells/mouse) from naive PLP TCR transgenic SJL mice were injected (i.v.) into CD11c-DTR *Rag2*−/− SJL BM chimeric mice (Day 0). Recipient mice had been treated on consecutive days before and after T cell transfer with DT (Sigma-Aldrich) or PBS (Day −1, 2, 4, 6, 8, 10, 12, 14).

For PLP-induced EAE, CD11c-DTR SJL BM chimeric mice that had been treated with DT (5 ng/g body weight) or PBS on day −3, −1, 1, and every other day for 30 days were injected (s.c.) with 75 μg PLP139-151 in CFA (Difco) into each flank. On the same day and 48 hrs later, each mouse received (i.v.) 200 ng pertussis toxin (PT, List Biological Laboratories). Subsequently, the mice were examined daily for clinical signs of EAE for 30 days.

For assessment of spontaneous EAE, PLP TCR Tg. CD11c-DTA or PLP TCR Tg. DTA control mice were examined at least twice a week for clinical signs of spontaneous EAE (EAE score ≥ 1) until four months of age. To assess passive or PLP-induced EAE, mice were examined daily for clinical signs of disease for indicated days. EAE severity was scored according to the following scale: 0, healthy; 1, limp tail; 2, partial hind limb paralysis; 3, complete hind limb paralysis; 4, complete hind and forelimb limb paralysis; 5, moribund.

### Statistical analysis

Differences in EAE incidence and severity were analyzed for statistical significance by Kaplan-Meier analysis/log-rank test or Fisher exact test and two-tailed Mann-Whitney U test, respectively. All other data were analyzed for statistical significance by Student’s *t*-test (two-tailed, unpaired). P values < 0.05 were considered statistically significant.

## Results

### Encephalitogenic T cells develop in PLP TCR Tg. CD11c-DTA SJL mice

Dendritic cells that present autoantigens promote negative selection of self-reactive thymocytes by inducing T cell apoptosis [29, 30]. Accordingly, T cell–mediated autoimmune diseases, including MS, have been associated with impaired or inefficient negative selection of autoreactive T cells in the thymus. We previously showed that myelin-specific T cells develop and are efficiently selected into peripheral lymphoid organs, resulting in spontaneous EAE in PLP TCR transgenic SJL mice [28]. To assess the requirement of DCs in the development and selection of PLP-specific CD4^+^ T cells, here we deleted CD11c^+^ DCs constitutively in PLP TCR transgenic SJL mice by Cre-lox technology.

CD11c-Cre C57BL/6 mice (B6.Cg-Tg(Itgax-cre)1-1Reiz/J) contain the Itgax-cre BAC transgene that drives the expression of Cre recombinase under the control of the integrin alpha × gene (*Itgax*, Cd11c) promoter/enhancer regions [26]. Rosa26-eGFP-DTA C57BL/6 mice (*Gt(ROSA)26Sor^tm1(DTA)Jpmb^*/J) harbor transgenes coding for *loxP*-flanked EGFP and diphtheria toxin A subunit (DTA). When bred to CD11c-Cre mice, DTA expression is activated by Cre-mediated excision of the *loxP*-flanked EGFP, which leads to specific ablation of Cre-expressing CD11c^+^ cells [27]. We fully backcrossed CD11c-Cre and Rosa26-eGFP-DTA transgenic C57BL/6 mice onto the SJL background and bred the offspring with PLP TCR transgenic SJL mice to generate PLP TCR Tg. CD11c-DTA SJL mice. In these novel mice, the gene for diphtheria toxin alpha chain (DTA) is activated selectively in CD11c^+^ DCs, resulting in constitutive DC ablation from birth. Thus, these mice allowed determining the requirement of DCs for thymic development and selection of PLP-specific CD4^+^ T cells. Notably, the frequency of CD11c^hi^ MHC II^+^ cDCs and CD11c^+^ CD11b^−^ B220^+^ pDCs [31] were reduced in the thymus of PLP TCR Tg. CD11c-DTA mice by 31- and 4-fold, respectively, as compared to PLP TCR Tg. DTA control mice. In the spleen, PLP TCR Tg. CD11c-DTA mice showed a >10-fold reduction in cDCs or pDCs as compared to their DC sufficient counterparts (Fig. 1 A). Similarly, we found profound reductions in the number of splenic cDCs (20-fold, p = 0.03) and pDCs (6-fold) in PLP TCR Tg. CD11c-DTA mice (Fig. 1 B).

**Figure 1.**
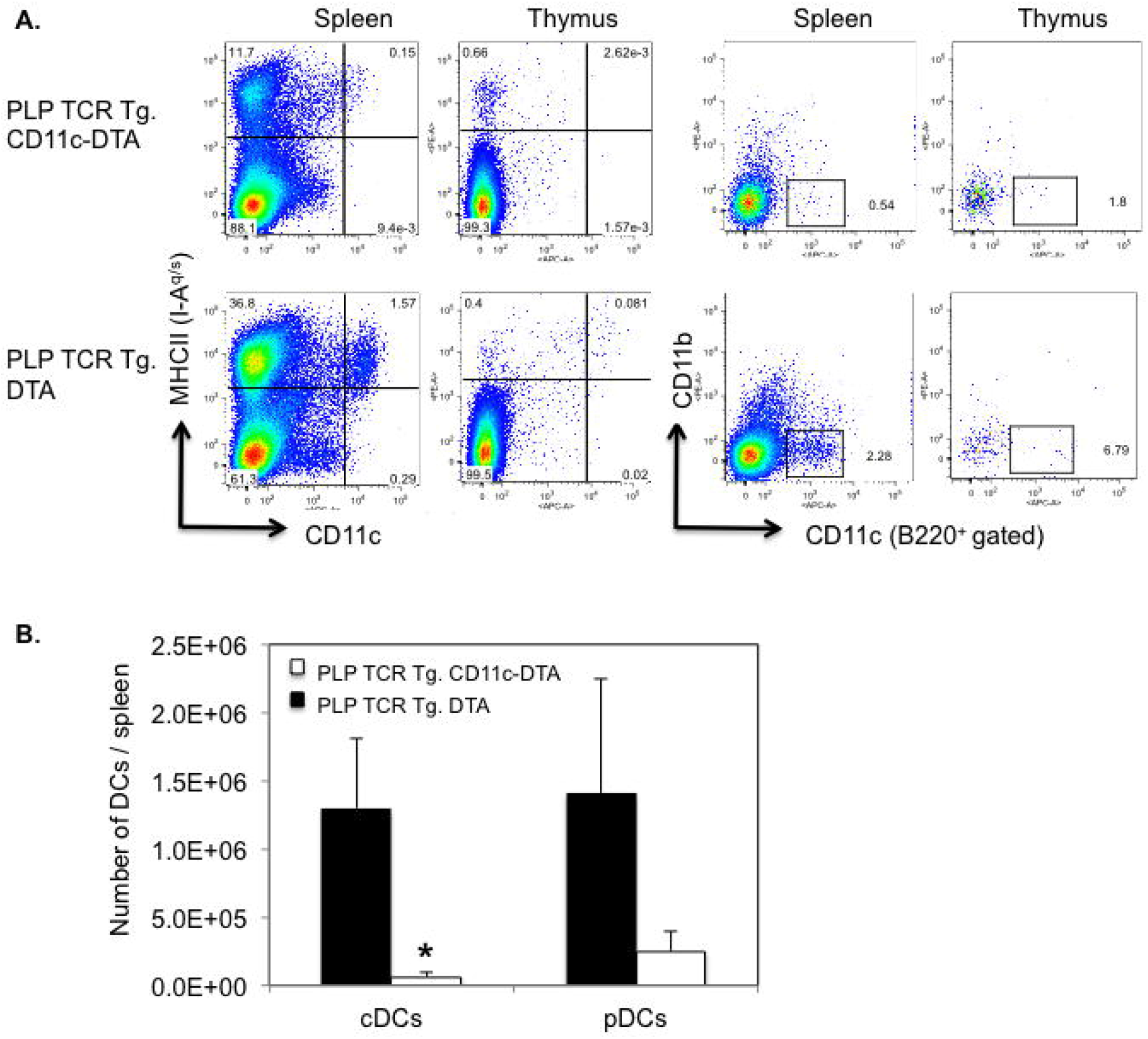
Efficient ablation of DCs in PLP TCR Tg. CD11c-DTA SJL mice. (A) Representative frequencies and (B) mean numbers ± SEM of cDCs (CD11c^hi^ MHC II^+^) and pDCs (CD11c^lo^ B220^+^ CD11b^−^) in the thymus (A) and spleen (A, B) of PLP TCR Tg. CD11c-DTA SJL mice (n = 4) or PLP TCR Tg. DTA SJL control mice (n = 3), as determined by flow cytometry, are shown. Numbers in dot plots represent percentages of live cells. Data shown are from four independent experiments. * p = 0.03 (Student’s t test)

DC-deficient and DC-sufficient PLP TCR transgenic mice showed comparable frequencies of thymic and peripheral T cell subsets. Furthermore, the profound deficiency in thymic DCs in PLP TCR Tg. CD11c-DTA mice did not appear to affect development or selection of CD4 and CD8 single positive nor double positive thymocytes because the their frequencies were similar to those observed in PLP TCR Tg. DTA mice (Fig. 2 A). As expected for MHC II restricted TCR transgenic mice, the peripheral T cell compartment in DC deficient and sufficient PLP TCR transgenic mice was skewed towards CD4^+^ T cells at the expense of CD8^+^ T cells (Fig. 2 A). Importantly, constitutive ablation of CD11c^+^ DCs did not result in significant differences in the number of total spleen cells nor in the number of T cell subsets or TCR transgenic (CD4^+^ TCR Vβ6^+^) T cells in PLP TCR Tg. CD11c-DTA as compared to DC sufficient control mice (Fig. 2 B).

**Figure 2.**
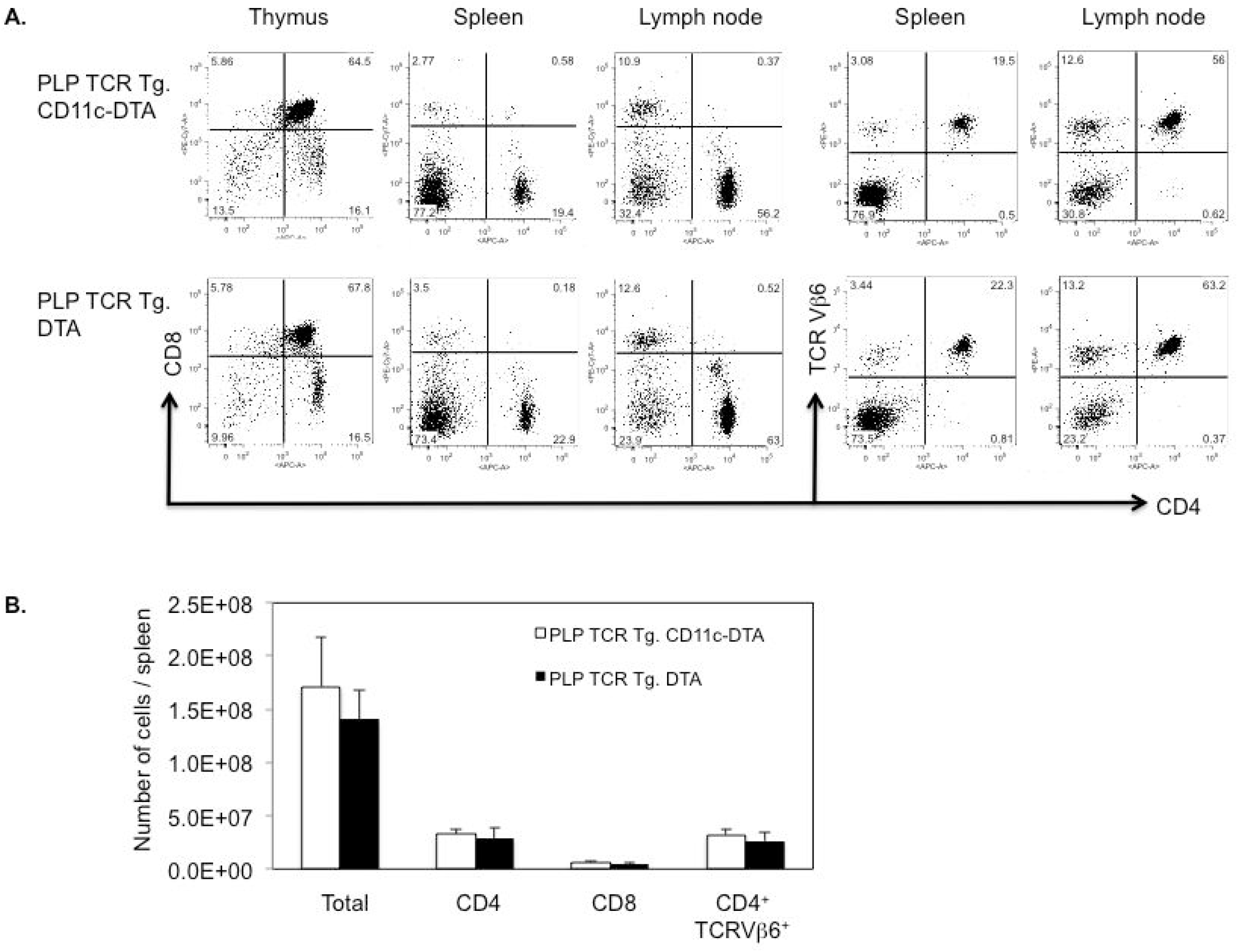
DC deficiency in PLP TCR Tg. CD11c-DTA SJL mice does not affect the development and selection of transgenic T cells. (A) Representative frequencies of T cell subsets in indicated lymphoid organs and (B) absolute numbers of total cells or indicated T cell subsets in the spleen of PLP TCR Tg. CD11c-DTA SJL mice (n = 4) or PLP TCR Tg. DTA control mice (n = 3), as assessed by flow cytometry, are shown. Numbers in dot plots represent percentages of cells. Data in bar graphs are represented as mean ± SEM. Data shown are from four independent experiments.

To assess the encephalitogenic function of PLP-specific Th cells that had developed in the absence of C11c^+^ DCs, we transferred purified CD4^+^ T cells from naïve PLP TCR Tg. CD11c-DTA SJL mice or PLP TCR Tg. DTA SJL mice, as controls, into RAG2 deficient SJL mice. PLP-specific CD4^+^ T cells from both transgenic lines mediated severe EAE in recipient mice within two weeks following T cell transfer (Fig. 3). Although CD4^+^ T cells from PLP TCR Tg. CD11c-DTA donor mice showed a tendency to mediate earlier and more severe signs of clinical EAE than those from PLP TCR Tg. DTA control mice, the differences did not reach statistical significance.

**Figure 3.**
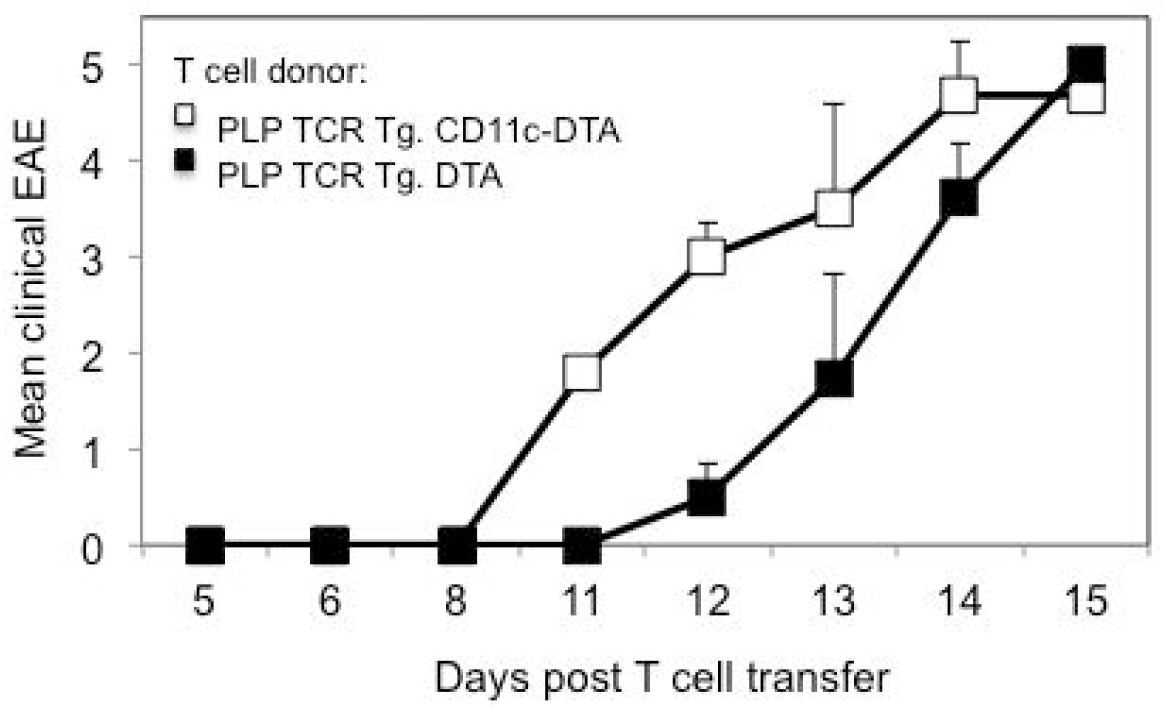
CD4^+^ T cells from PLP TCR Tg. CD11c-DTA SJL mice mediate severe EAE in *Rag2−/−* SJL recipient mice. Column purified CD4^+^ T cells from naïve PLP TCR Tg. CD11c-DTA SJL mice or PLP TCR Tg. DTA control mice were adoptively transferred into *Rag2 −/−* SJL recipient mice (n = 4-5 mice/group). Clinical signs of EAE were assessed daily in recipient mice for the indicate time and are shown as mean clinical EAE ± SEM.

In summary, these data demonstrate that the profound reduction of DCs in PLP TCR Tg. CD11c-DTA mice did not affect the thymic development and selection of autoreactive PLP TCR transgenic CD4^+^ T cells. Additionally, CD4^+^ T cells from naïve PLP TCR Tg. CD11c-DTA mice were not impaired in their capacity to mediate severe EAE in recipient mice.

### PLP TCR Tg. CD11c-DTA SJL mice develop accelerated spontaneous EAE

We and other investigators previously reported high incidence of spontaneous EAE in PLP-specific TCR transgenic SJL mice, housed at different animal facilities [28, 32, 33]. The requirement and the role of DCs in the pathogenesis of spontaneous EAE are unknown.

To address this gap in knowledge, we examined in PLP TCR Tg. CD11c-DTA SJL mice and PLP TCR Tg. DTA SJL mice as controls at least twice per week for clinical signs of spontaneous EAE (EAE score ≥ 1) for up to 120 days after birth. While all monitored mice of both transgenic lines developed spontaneous EAE (100% incidence), disease onset time was significantly different between the two lines. The median (35 vs. 78 days; 95% CI = 6.953E-310 – 6.084E-310; p = 0.0005, log-rank test) and mean day of EAE onset (39.7 ± 4.3 vs. 80.6 ± 6.3 days ± SEM; p < 0.0001, two-tailed Mann-Whitney test) in PLP TCR Tg. CD11c-DTA SJL mice were significantly earlier than in PLP TCR Tg. DTA SJL control mice (Fig. 4 and Table 1). The mean maximum EAE severity was increased in PLP TCR Tg. CD11c-DTA SJL mice as compared to the DC sufficient control mice (3.4 ± 0.5 vs. 2.9 ± 0.3, mean EAE score ± SEM; p = 0.1, two-tailed Mann-Whitney U test) but did not reach statistical significance (Fig. 4 and Table 1).

**Figure 4.**
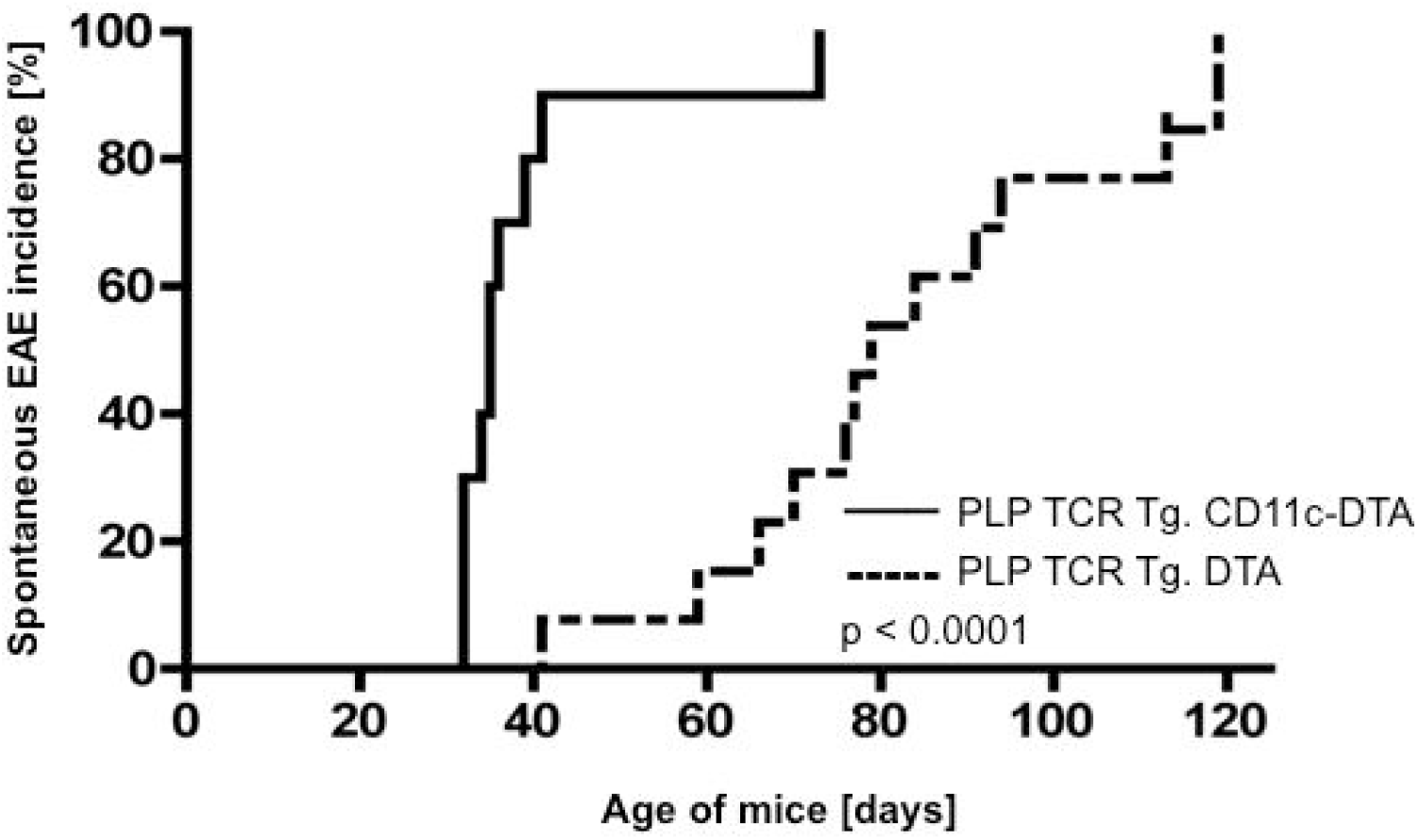
Accelerated onset of spontaneous EAE in PLP TCR Tg. CD11c-DTA SJL mice. PLP TCR Tg. CD11c-DTA SJL mice (n = 9) or PLP TCR Tg. DTA mice (n = 12) were examined twice per week for clinical signs of spontaneous EAE (EAE score ≥ 1) during the indicated age of mice. Incidence of spontaneous EAE and statistical significance were determined by Kaplan–Meier analysis and log rank test, respectively. p < 0.0001

**Table 1.** Spontaneous EAE in DC deficient PLP TCR transgenic CD11c-DTA SJL mice.

| Mice | EAE incidence [%] | Mean age of EAE onset ± SEM [Days] | Mean max. EAE severity ± SEM |
| --- | --- | --- | --- |
| PLP TCR Tg. CD11c-DTA (n = 9) | 100 | 39.7 ± 4.3 | 3.4 ± 0.5 |
| PLP TCR Tg. DTA (n = 12) | 100 | 80.6 ± 6.3 | 2.9 ± 0.3 |
| Significance | $p > 0.05$ <sup>a</sup> | $p = 0.0005$ <sup>b</sup> | $p > 0.05$ <sup>c</sup> |

<sup>a</sup> Fisher exact test
<sup>b, c</sup> Two-tailed Mann-Whitney U test

Earlier onset of spontaneous EAE in PLP TCR Tg. CD11c-DTA SJL mice was associated with significantly higher frequencies of splenic CD4^+^ T cells, expressing the early T cell activation marker CD69 as compared to PLP TCR Tg. DTA SJL (16.0% ± 1.8% vs. 5.2% ± 0.6%; p = 0.004). Additionally, PLP TCR Tg. CD11c-DTA mice harbored higher frequencies of effector memory CD4^+^ T cells than the control mice as judged by the percentages of CD62L^lo^ splenocytes (48.8% ± 4.1% vs. 35.5% ± 3.4%; mean ± SEM, p > 0.05), although this difference was not statistically significant (Fig. 5).

**Figure 5.**
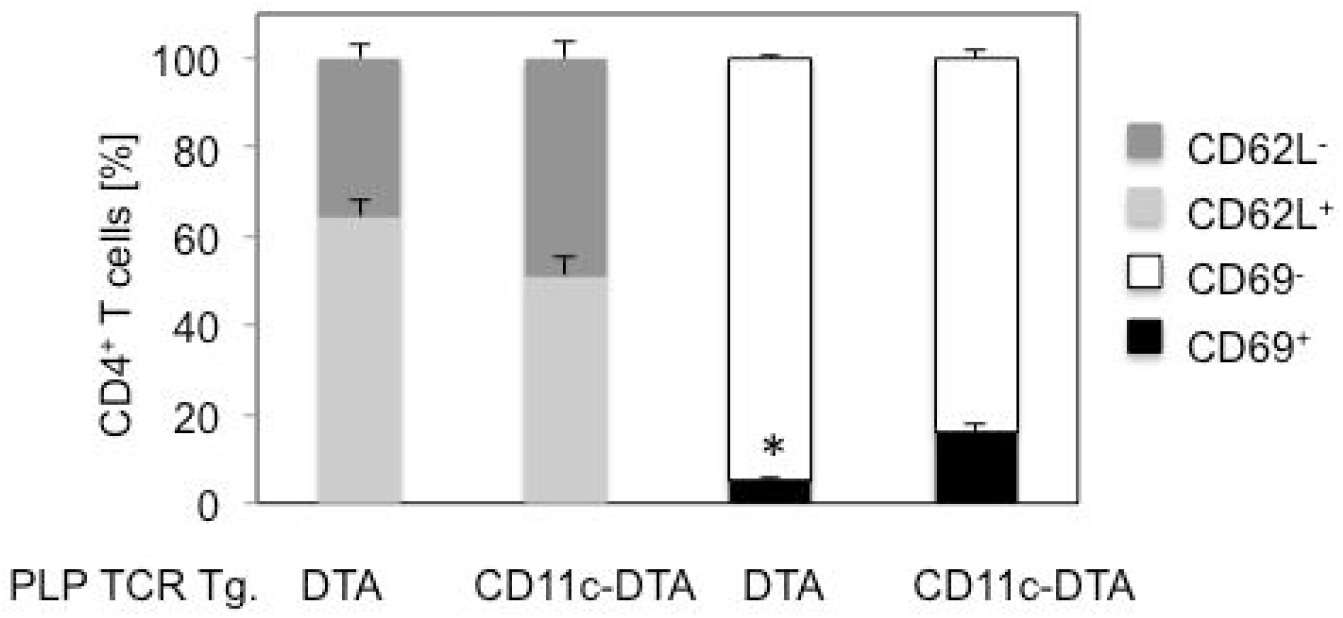
Increased frequencies of primed CD4^+^ T cells in PLP TCR Tg. CD11c-DTA SJL mice. Frequencies of early activated (CD69^+^), effector memory (CD62L^−^) cells or unprimed cells (CD69^−^, CD62L^+^) in splenic CD4^+^ T cells from 5-8 week old PLP TCR transgenic CD11c-DTA SJL mice (n = 4) or PLP TCR Tg. DTA control mice (n = 3) were determined by flow cytometry. Data are shown as mean cell frequencies (%) ± SEM of four independent experiments. * p = 0.004 (Student’s t test)

Taken together, these results show that constitutive ablation of DCs in PLP TCR Tg. CD11c-DTA SJL significantly accelerated the development of spontaneous EAE, which was associated with enhanced activation and priming of naïve CD4^+^ T cells.

### Profound depletion of DCs in DT treated CD11c-DTR SJL mice

CD11c-DTR C57BL/6 mice (B6.FVB-*1700016L21Rik^Tg(Itgax-DTR/EGFP)57Lan^*/J) contain a transgene that encodes the simian diphtheria toxin receptor fused to enhanced green fluorescent protein (DTR/EGFP) under the control of the murine CD11c promoter, which directs DTR/EGFP expression to dendritic cell populations [12]. To investigate the requirement of CD11c^+^ DCs in priming PLP-specific CD4^+^ T cells and EAE induction in non-TCR transgenic mice, we established novel CD11c-DTR SJL mice by fully backcrossing CD11c-DTR C57BL/6 mice [12] onto the autoimmune prone SJL/J strain. CD11c-DTR SJL mice allowed selective conditional ablation of CD11c^+^ DCs upon diphtheria toxin (DT) administration due to the CD11c promoter driven expression of a transgene encoding the human DT receptor (DTR)-GFP fusion protein.

We determined the efficiency of DT induced depletion of DCs by assessing the frequencies and absolute numbers of splenic cDCs and pDCs in CD11c-DTR SJL mice, following DT injections for two consecutive days. Flow cytometric analysis showed a 10-fold reduction in the frequency of DTR-GFP transgenic (CD11c^hi^ GFP^+^) DCs or CD11c^hi^ MHC II (I-A^q^)^+^ cDCs (p = 0.01) in DT treated as compared to PBS treated CD11c-DTR SJL mice (Fig. 6 A). Absolute numbers of splenic cDCs were ≥ 10-fold lower (p = 0.02) in DT treated than in PBS treated CD11c-DTR SJL mice (Fig. 6 B). The frequency and absolute numbers of splenic pDCs, defined as CD11c^lo^ B220^+^ CD11b^−^ cells, were reduced by 2- and 4-fold, respectively in DT treated CD11c-DTR SJL mice as compared to control treated mice, which was statistically not significant (p > 0.05) (Fig. 6 A and 6 B). Thus, two consecutive injections of DT were sufficient to achieve profound ablation of cDC and detectable depletion of pDCs in CD11c-DTR SJL mice, which was also reported for cDCs but not for pDCs in non-autoimmune prone strains of mice [12, 24].

**Figure 6.**
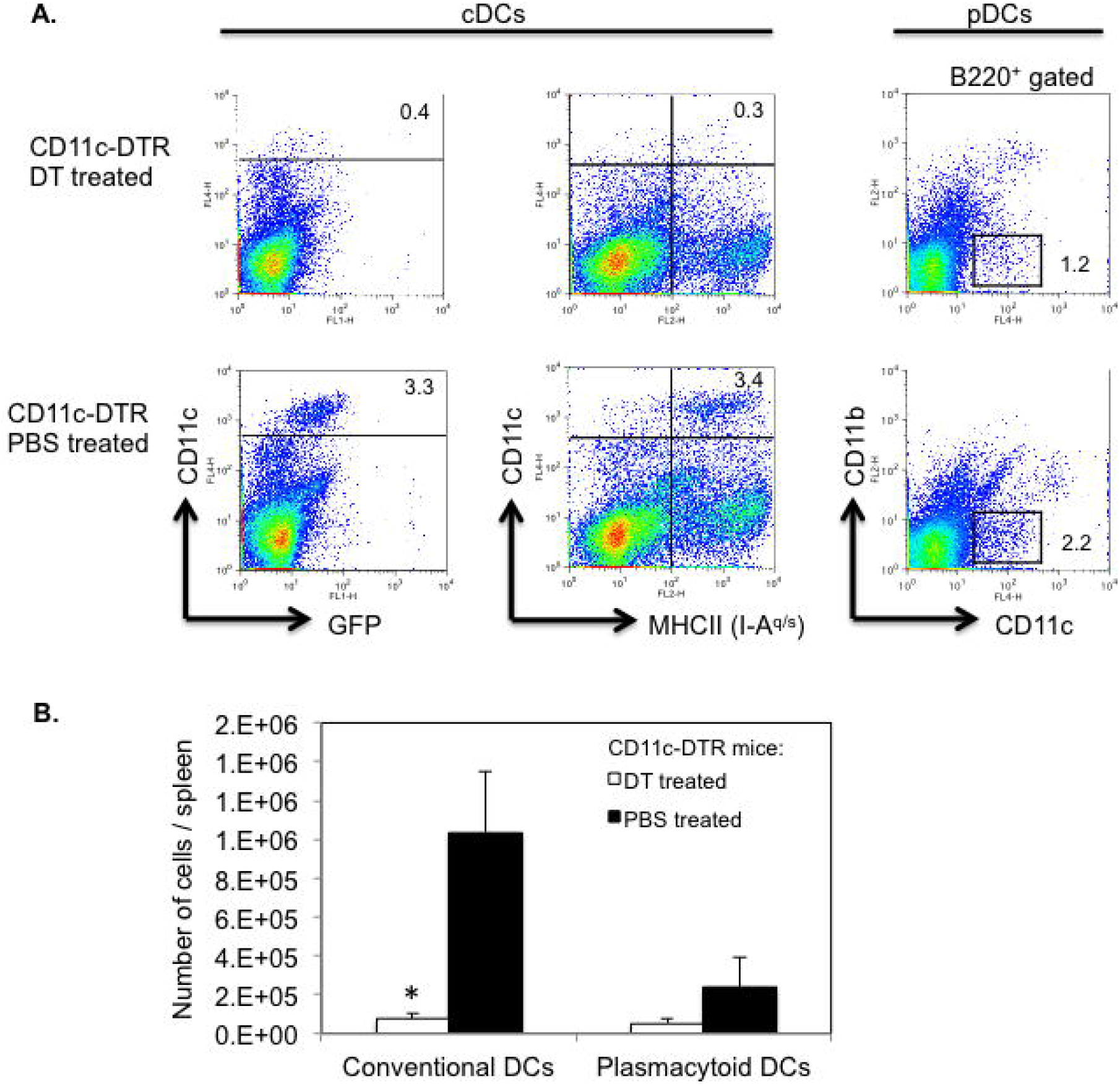
Efficient depletion of DCs in DT treated CD11c-DTR SJL mice. CD11c-DTR SJL mice were injected on two consecutive days with DT or PBS (2 - 3 mice/group). (A) Following the final injection, spleen cells from mice were isolated and assessed for cDCs (CD11c^hi^ MHC II^+^ or CD11c^hi^ GFP^+^) or pDCs (CD11c^lo^ B220^+^ CD11b^−^) by flow cytometry. Representative data of two independent experiments are shown. Numbers in dot plots refer to percentages of DC subpopulations. (B) Absolute numbers of cDCs (CD11c^hi^ MHC II^+^) and pDCs (CD11c^lo^ B220^+^ CD11b^−^) in DT treated or PBS treated mice. Data are shown as mean number of cells/spleen ± SEM from two independent experiments. * p < 0.02 (Student’s t test)

### Efficient priming and differentiation of PLP-specific CD4^+^ T cells in DC depleted SJL mice

Next, we investigated the requirement of DCs for priming and differentiation of myelin-specific CD4^+^ T cells in vivo by assessing antigen-specific proliferation and Th cytokine responses in PLP-specific CD4^+^ T cells following transfer into DT treated CD11c-DTR SJL mice. CD4^+^ T cells from naïve PLP TCR transgenic SJL mice were labeled with CellTrace Violet and transferred (day 0) into DT treated (day −1, 1, 3) CD11c-DTR SJL mice or DC sufficient control mice. On the day of the T cell transfer, the mice were injected subcutaneously with PLP139-151 in CFA. Four days later (day 4), we isolated cells from draining lymph nodes (axillary, brachial) of the mice and assessed the dye dilution of labeled PLP TCR transgenic (CD4^+^ TCR Vβ6^+^) T cells by flow cytometry to determine their proliferative response to cognate antigen. Transgenic T cells from DC depleted or DC sufficient recipient mice had undergone up to 7 cell divisions, indicating vigorous PLP-specific T cell proliferation in both groups (Fig. 7 A). The frequencies of divided CD4^+^ TCR Vβ6^+^ T cells in DT treated CD11c-DTR SJL recipient mice were similar to those in DC sufficient control recipients (76.1% ± 3.4% vs. 78.1% ± 2.4%; mean ± SEM) (Fig. 7 B).

**Figure 7.**
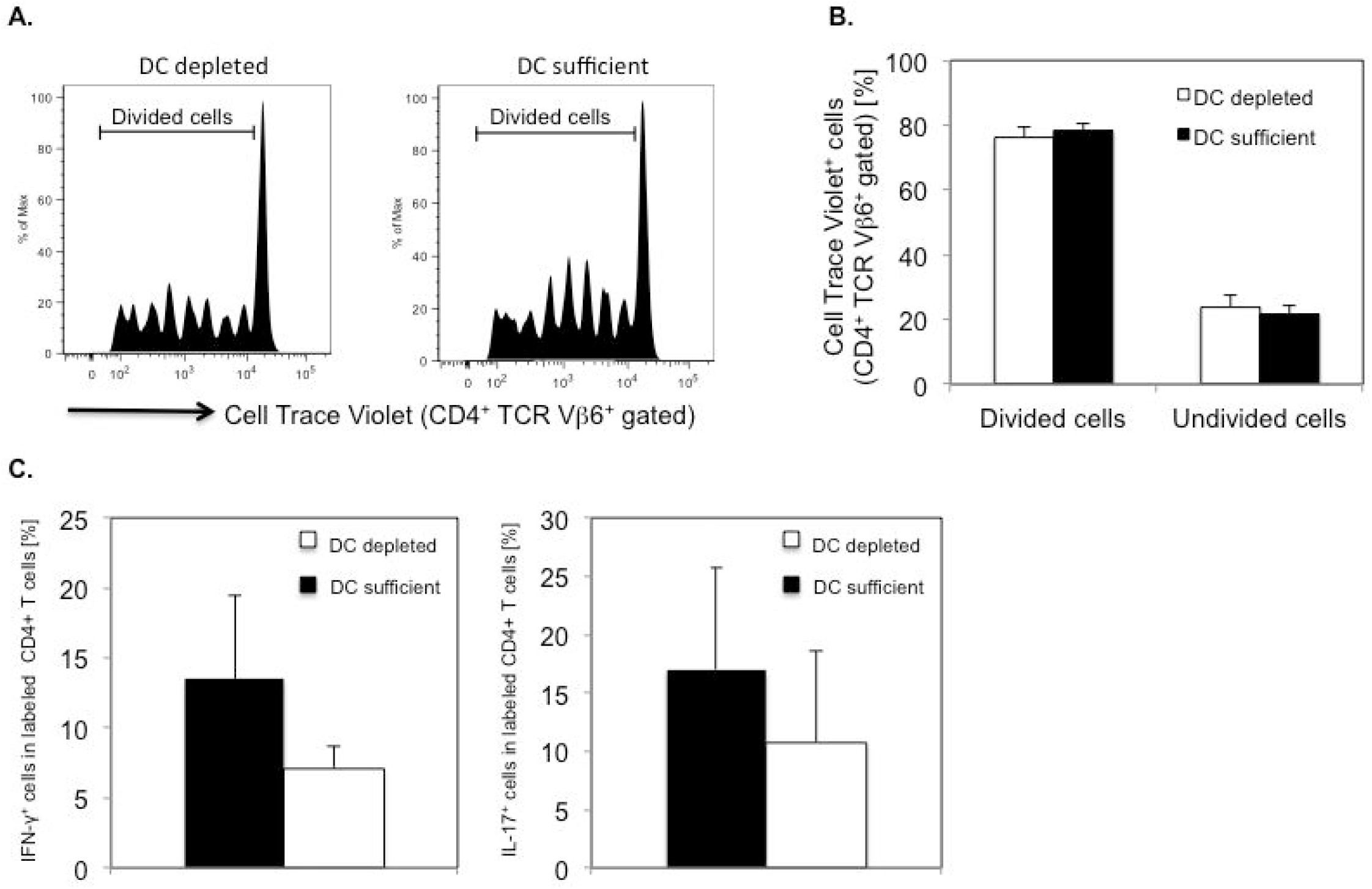
Efficient PLP-specific proliferation and cytokine responses by PLP TCR transgenic CD4^+^ T cells in DT treated CD11c-DTR SJL mice. DT treated CD11c-DTR SJL mice (n = 5) or DC sufficient control mice (n = 4) received CellTrace Violet labeled CD4^+^ T cells from naïve PLP TCR transgenic SJL mice. The mice were subsequently injected (s.c.) with PLP139-151 in CFA. Four days later, draining lymph node cells from one mouse of each group, were analyzed for cell divisions in PLP TCR transgenic (CD4^+^ TCR Vβ6^+^ gated) T cells by flow cytometry. (A) Representative cell divisions in CD4^+^ TCR Vβ6^+^ gated cells and (B) Mean frequencies ± SEM (%) of undivided (0 cell division) and divided (> 1 cell division) CD4^+^ TCR Vβ6^+^ gated cells from recipient mice of four independent experiments are shown. (C) Draining lymph node cells from DT treated CD11c-DTR SJL mice or control mice (n = 1 mouse/group) were stimulated overnight with PLP139-151 in vitro and assessed for intracellular cytokine production by flow cytometry. Data are shown as mean frequencies ± SEM (%) of CellTrace Violet^+^ CD4^+^ T cells expressing IFN-γ or IL-17 of three independent experiments.

To investigate whether DCs were required for promoting PLP-specific CD4^+^ T cell differentiation, we assessed effector Th cytokines in isolated lymph node cells, following overnight stimulation with PLP-139-151 in vitro. Labeled CD4^+^ T cells from DT treated CD11c-DTR SJL recipients showed lower frequencies of IFN-γ producing Th1 cells or IL-17 producing Th17 cells than those from DC sufficient recipients, as determined by intracellular cytokine analysis (IFN-γ: 7.1% ± 1.6% vs. 13.5% ± 6.0%; IL-17: 10.8% ± 7.9% vs. 17.0% ± 8.7%; mean ± SEM). Notably, the frequencies of cytokine expressing CD4^+^ T cells varied substantially among individual mice and were statistically not significantly different between the two groups of mice (Fig. 7 C). We did not observe detectable frequencies of CD4^+^ T cells that produced Th2 signature cytokines such as IL-4 or IL-10, following PLP-specific re-stimulation (data not shown).

Taken together, these data demonstrate that in the absence of DCs, naïve PLP TCR transgenic CD4^+^ T cells were able to proliferate and differentiate into Th1 and Th17 effector cells in response to cognate antigen stimulation in vivo. These findings suggest that cells other than DCs efficiently presented PLP antigen and primed CD4^+^ T cells in DT treated CD11c-DTR SJL mice.

### Conditionally DC depleted SJL mice are highly susceptibility to PLP-induced EAE

Having demonstrated that pro-inflammatory PLP-specific CD4^+^ T cell responses could be promoted in DC deficient mice, we next determined whether DCs were essential for inducing encephalitogenic T cell function and EAE.

As others previously reported, CD11c-DTR mice show increased incidences of mortality following repetitive injections of DT, which is presumably due to DT-induced deletion of DTR expressing non-hematopoietic cells [12, 34]. In contrast, mixed irradiation CD11c-DTR chimeras are not susceptible to DT induced mortality [34]. Thus, to allow long-term depletion of CD11c^+^ DCs, we generated mixed CD11c-DTR bone marrow (BM) chimeric mice by reconstituting lethally irradiated SJL hosts with BM from CD11c-DTR SJL mice. Eight weeks following BM transplantation, chimeras were injected with DT (5 ng/g body weight) or with PBS, as control on day −3 and every other day for the entire duration of the experiment. To induce EAE, chimeric mice were injected (day 0) with PLP139-151/CFA and pertussis toxin (Fig. 8 A). Notably, 100% of DT treated CD11c-DTR BM chimeric mice succumbed to EAE as compared to 50% of PBS treated control chimeras. DT and PBS treated CD11c-DTR BM chimeric mice that showed clinical signs of EAE had comparable disease onset time (13.0 ± 1.3 vs. 13.5 ± 2.5; mean day of onset ± SEM) and similar mean maximum severity (3.2 ± 0.7 vs. 3.5 ± 1.5; mean maximum EAE severity ± SEM) (Fig. 8 B and Table 2). The differences in these disease parameters between the two groups were statistically not significant.

**Figure 8.**
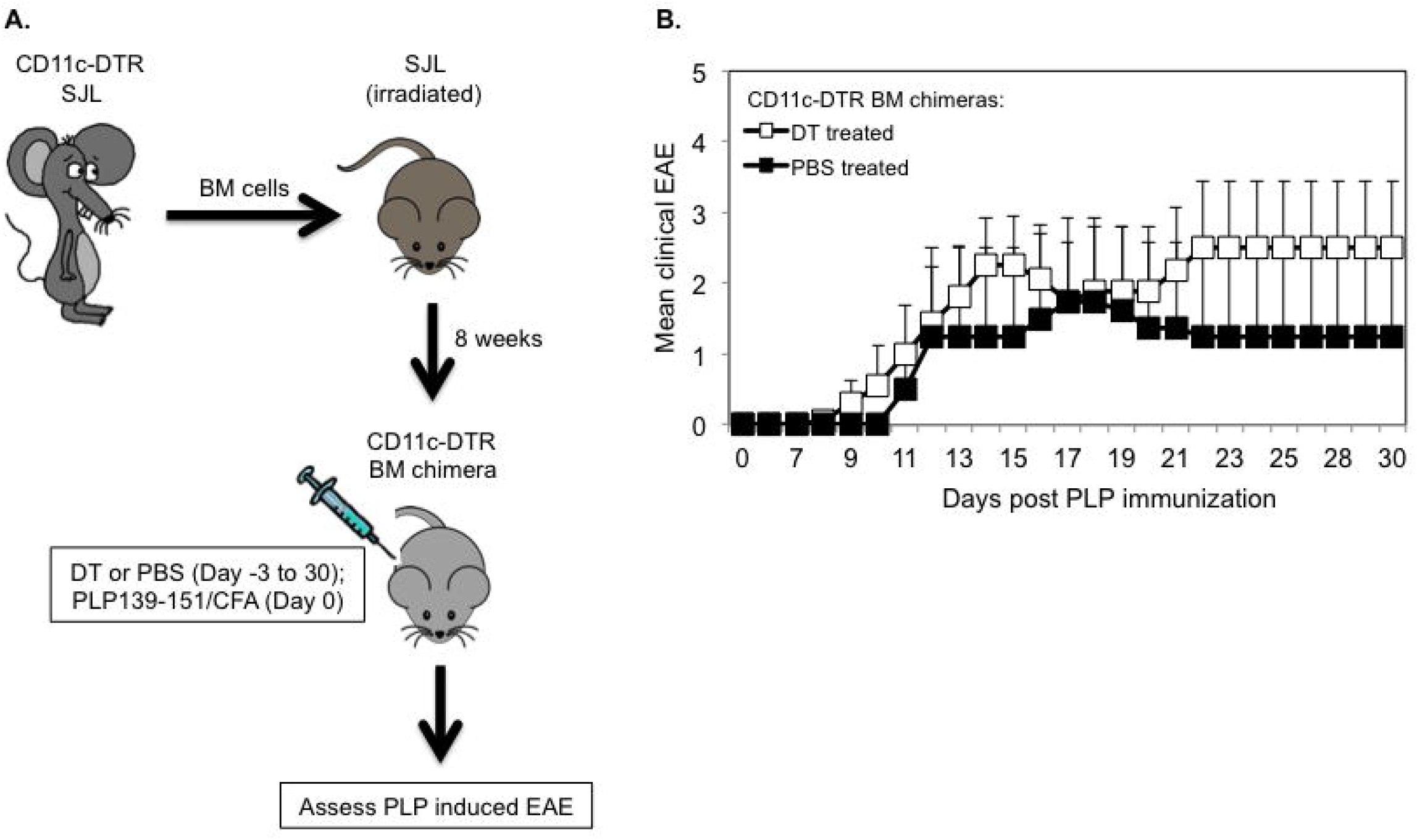
Long term DT treated CD11c-DTR SJL bone marrow chimeric mice are susceptible to PLP-induced EAE. (A)Schematic representation of the generation of CD11c-DTR SJL BM chimeric mice and assessment of PLP-induced EAE in long term DT or PBS treated chimeras. (B)EAE was induced in DT (n = 8) or PBS (n = 4) treated CD11c-DTR SJL BM chimeric mice by injection of PLP139-131/CFA (s.c.) and PT (i.v). The mice were examined daily for clinical signs of EAE for 30 days during which they received DT or PBS on every other day. Data are shown as mean clinical EAE ± SEM.

**Table 2.** PLP-induced EAE in DT treated CD11c-DTR SJL BM chimeras.

| **CD11c-DTR SJL**<br>**BM chimeras** | **EAE incidence [%]** | **Mean day of EAE**<br>**onset ± SEM** | **Mean max. EAE**<br>**severity ± SEM** |
| --- | --- | --- | --- |
| DT treated (n = 8) | 100 | $13.0 \pm 1.3$ | $3.2 \pm 0.7$ |
| PBS treated (n = 4) | 50 | $13.5 \pm 2.5$ | $3.5 \pm 1.5$ |
| Significance | $p > 0.05^a$ | $p > 0.05^b$ | $p > 0.05^b$ |

<sup>a</sup> Fisher exact test
<sup>b</sup> Two-tailed Mann-Whitney U test

These results demonstrate that DCs were dispensable for the priming of encephalitogenic T cells and EAE induction following PLP immunization in long term DT treated CD11c-DTR SJL BM chimeras.

### Transfer of PLP-specific CD4^+^ T cells accelerates EAE onset in long term DC depleted SJL mice

Purified CD4^+^ T cells from naïve PLP TCR transgenic SJL mice mediate severe EAE within two weeks following transfer into syngeneic RAG2 deficient recipient mice [32]. It is unclear whether APCs such as DCs presenting endogenous autoantigen are required in activating PLP-specific T cells resulting in passive EAE. To investigate this question, we reconstituted lethally irradiated RAG2 deficient SJL mice with BM cells from CD11c-DTR *Rag2* −/− SJL mice we had generated by crossing and intercrossing CD11c-DTR with RAG2 deficient SJL mice (Fig. 9 A). Eight weeks following BM reconstitution, we transferred purified CD4^+^ T cells from naïve PLP TCR transgenic SJL mice into mixed irradiation CD11c-DTR *Rag2* −/− SJL chimeras. The recipient mice were treated with DT (5 ng/g body weight) the previous day and every other day (day −1, 2, 4, 6, 8, 10, 12, 14) after T cell transfer for two weeks to ablate their DCs or, as control, injected with PBS. Similar to other reports, DT induced long-term ablation of DCs did not cause any mortality and was highly effective. Compared to PBS treated control mice, DT treated mice showed > 10-fold reduction in frequency (1% vs. < 0.1%) and significantly fewer numbers (285,200 DCs ± 104,972 vs. 27,975 ± 2,096; mean DC number ± SEM; p = 0.03) of splenic CD11c^+^MHC II^hi^ DCs (Fig. 9 B and 9 C).

**Figure 9.**
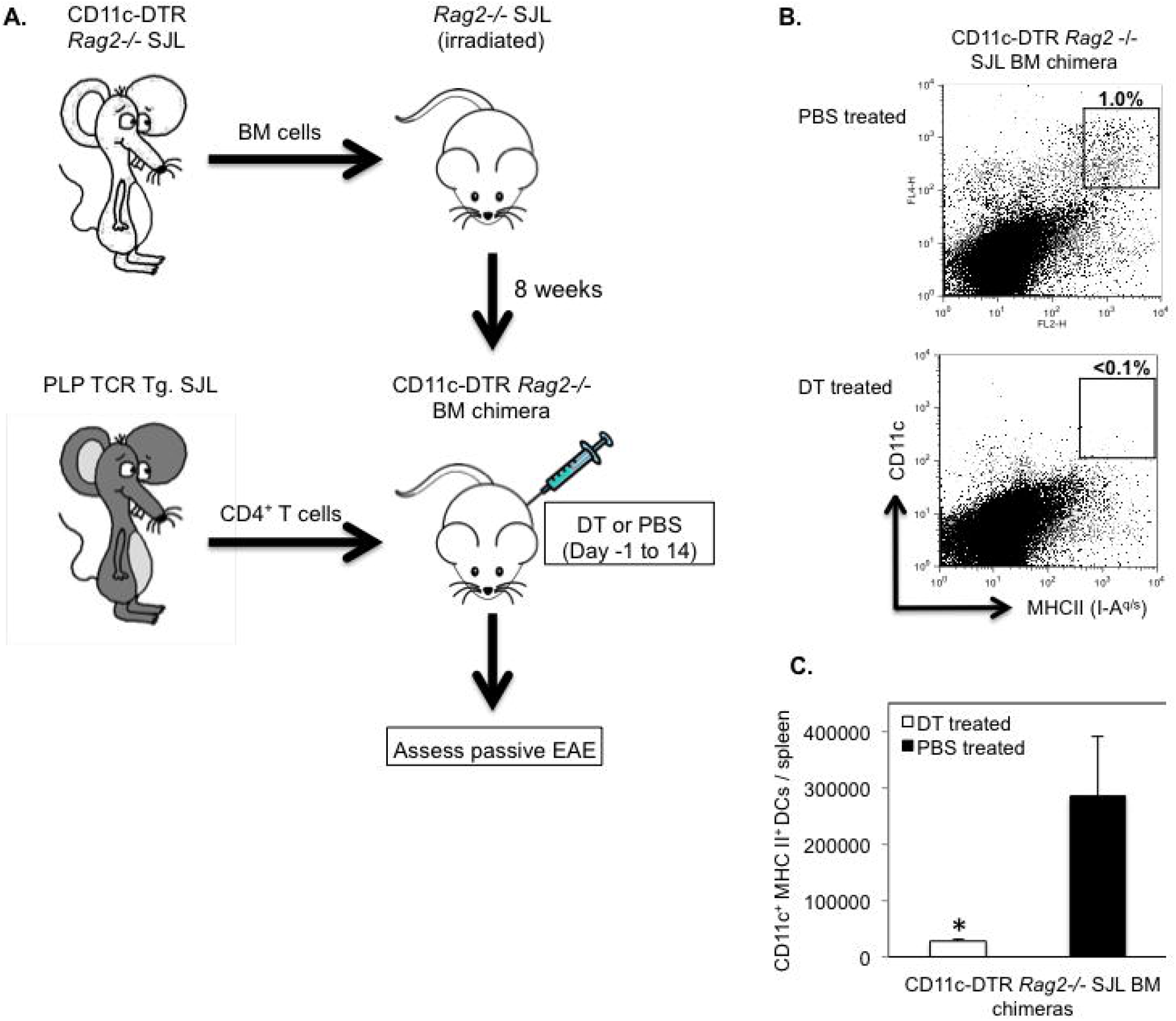
Efficient DC ablation in long term DT treated CD11c-DTR *Rag2*−/− SJL BM chimeras. (A) Schematic representation of the generation of CD11c-DTR *Rag2 −/−* SJL BM chimeric mice and assessment of their susceptibility to passive EAE by transfer of CD4^+^ T cells from naïve PLP TCR transgenic SJL mice. (B, C) Splenic DCs from CD11c-DTR *Rag2* −/− SJL BM chimeras that had been treated with DT (n = 4) or PBS (n = 3) on consecutive days for two weeks were assessed by flow cytometry at the end of the treatment. (B) Representative frequency (number above boxed gates, %) or (C) Mean absolute numbers ± SEM of CD11c^+^ MHC II (IA-^q^)^+^ DCs in live splenocytes from long term treated CD11c-DTR *Rag2* −/− SJL BM chimeras of three independent experiments are shown. * p < 0.03 (Student’s t test)

Notably, all DT treated or PBS treated (100%) CD11c-DTR *Rag2* −/− BM chimeras developed severe EAE resulting in a moribund state or death within two weeks following transfer of PLP-specific CD4^+^ T cells. The mean day of EAE onset was significantly earlier in DC depleted recipients than in control mice (10.0 ± 0.3 vs. 11.7 ± 0.5; mean day of EAE onset ± SEM; p < 0.05) (Table 3).

**Table 3.** Passive EAE in DT treated CD11c-DTR *Rag2*−/− SJL BM chimeras.

| **CD11c-DTR *Rag2*<sup>-/-</sup>**<br>**SJL BM chimeras** | **EAE incidence [%]** | **Mean day of EAE**<br>**onset ± SEM** | **Mean max. EAE**<br>**severity ± SEM** |
| --- | --- | --- | --- |
| DT treated (n = 8) | 100 | 10 ± 0.3 | 5 ± 0 |
| PBS treated (n = 9) | 100 | 11.7 ± 0.5 | 5 ± 0 |
| Significance | p > 0.05<sup>a</sup> | p = 0.03<sup>b</sup> | p > 0.05<sup>b</sup> |

<sup>a</sup> Fisher exact test<sup>b</sup> Two-tailed Mann-Whitney U test

The mean severity of EAE was significantly increased in DT treated recipients as compared to PBS treated controls during the early induction phase of the disease (Day 10: 1.5 ± 0.5 vs. 0.4 ± 0.3 mean maximum EAE severity ± SEM; p < 0.03. Day 11: 3.2 ± 0.5 vs. 1.6 ± 0.6; mean maximum EAE score ± SEM; p < 0.05) (Fig. 10 A). However, the mean maximum EAE severity throughout the study was not significantly different between the two treatment groups (Table 3). Consistent with efficient activation of naive PLP-specific T cells, the majority of TCR transgenic CD4^+^ T cells in DT treated recipient mice affected with severe EAE (EAE score ≥ 4) expressed CD44 (72.8% ± 4.1%; mean frequency ± SEM), a marker of activated/memory T cells (Fig. 10 C). The frequency of the same CD4^+^ T cell subpopulation was slightly increased in diseased control mice (81.9% ± 9%; mean frequency ± SEM) but statistically not different to that in DT treated recipient mice (p > 0.05).

**Figure 10.**
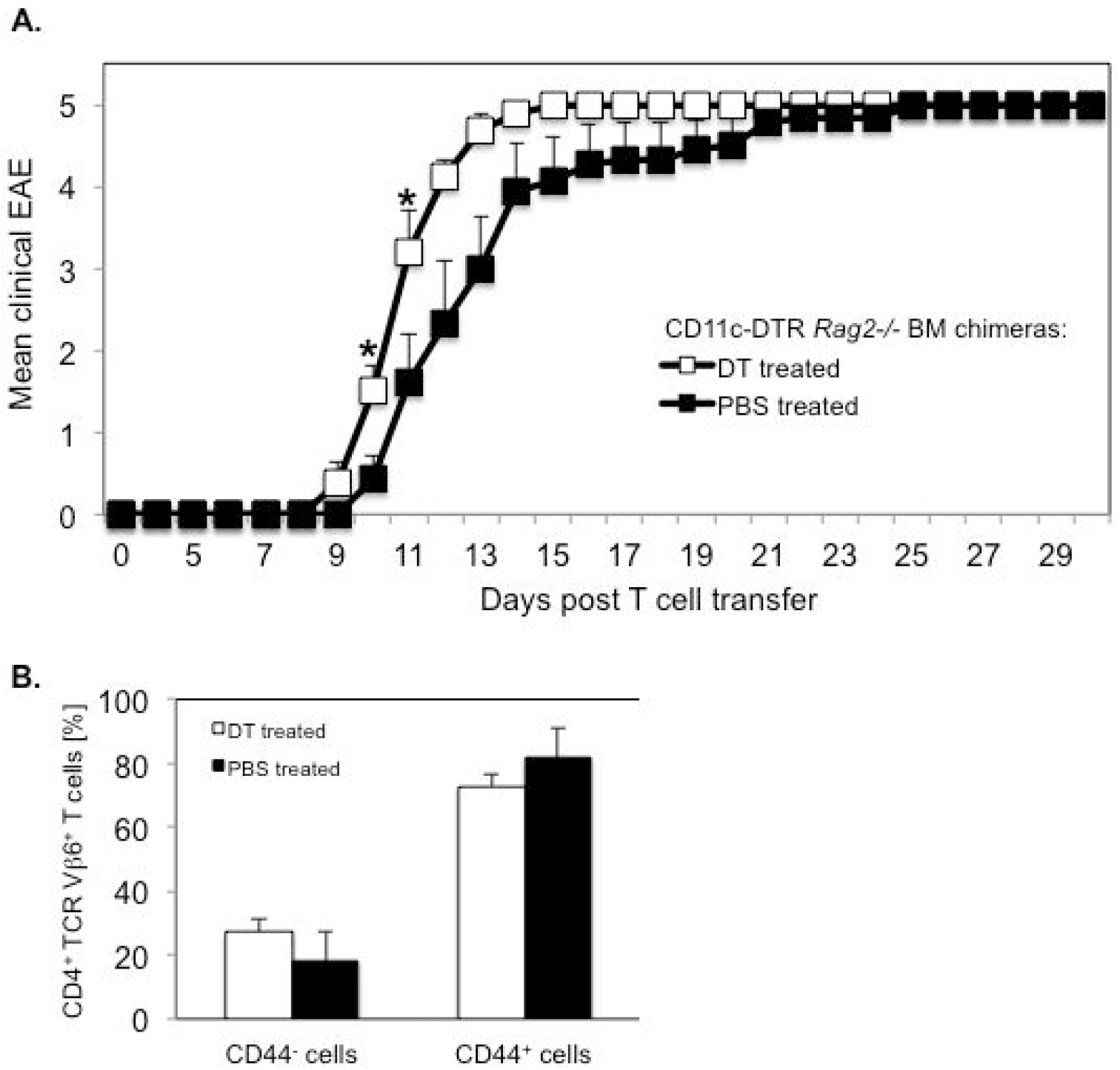
Transfer of PLP-specific CD4^+^ T cells induces severe EAE in long term DT treated CD11c-DTR *Rag2*−/− SJL BM chimeras. CD11c-DTR *Rag2*−/− SJL BM chimeric mice were treated with DT or PBS on consecutive days before and following the transfer of CD4^+^ T cells from naïve PLP TCR transgenic SJL mice for two weeks. Recipient mice were examined for clinical signs of EAE for 30 days. (A) Mean clinical EAE + SEM in DT treated (n = 8) or PBS treated (n=9) recipient mice from two independent experiments is shown. (B) Spleen cells from DT treated (n = 5) or PBS treated (n = 3) mice affected with severe EAE (EAE score ≥ 4) were analyzed for expression of the CD44 activation marker in transgenic T cells by flow cytometry. Frequencies of CD44^+^ (activated / memory) or CD44^−^ (non-activated) cells in transgenic (CD4^+^ TCR Vβ6^+^ gated) T cells are shown as mean percentages ± SEM.

Collectively, these results demonstrate that naïve PLP-specific CD4^+^ T cells were primed by endogenous autoantigen even in the absence of DCs in DT treated CD11c-DTR R*ag2* −/− BM chimeric mice, resulting in accelerated onset of severe EAE. These findings indicate that DCs were not required for presenting autoantigen to PLP-specific CD4^+^ T cells, and that non-DCs compensated for the lack of DCs by efficiently promoting encephalitogenic T cell responses. The data further show that DC ablation in DT treated CD11c-DTR R*ag2* −/− BM chimeric mice led to an acceleration of EAE onset, which was conceivably due to the loss of tolerogenic DCs.

## Discussion

DCs are considered the most potent APCs to elicit T cell responses against infectious agents [35]. To define cellular mechanisms that regulate the development of autoimmunity, the role of DCs have been studied in different autoimmune disease models, including EAE [36]. Studies in EAE involved in vitro derived DCs or DC subsets isolated from mice, which requires extensive experimental manipulations that can affect the phenotype and function of the examined DCs [18, 37, 38]. More recently, genetically engineered mouse models allowing selective depletion of DCs in vivo, have been utilized to examine the role of DCs in MOG-induced EAE in non-autoimmune prone C57BL/6 mice [24, 25]. Although these studies were informative, they could not study the potential contribution of DCs in myelin-specific T cell responses in a spontaneous autoimmune disease setting. In addition, those studies assessed the requirement of DCs in EAE mediated by T cells exclusively reactive to MOG, a minor myelin antigen, but not to one of the major CNS antigens such as PLP. To address this knowledge gap, we established PLP TCR Tg. CD11c-DTA and CD11c-DTR mice on the SJL strain, which is highly susceptible to EAE mediated by CD4^+^ T cells specific for PLP, a potential autoantigen in MS [39, 40]. These novel mice allowed constitutive or conditional depletion of CD11c^+^ DCs to investigate their requirement in spontaneous EAE and EAE induced by PLP immunization or by adoptive transfer of PLP-specific CD4^+^ T cells.

We show that genetic ablation of DCs from birth in PLP TCR Tg. CD11c-DTA SJL mice depleted cDCs almost completely and pDCs substantially in the thymus, as well as the peripheral lymphoid organs of these mice. Constitutive loss of DCs did not affect thymic or peripheral selection of self-reactive T cells in PLP TCR Tg. CD11c-DTA SJL mice because we detected comparable frequencies of CD4^+^ T cells transgenic for the PLP-specific TCR Vβ6 chain in peripheral lymphoid organs of DC deficient or DC sufficient TCR transgenic mice. This finding is consistent with results obtained in DC depleted non-TCR transgenic mice [41, 42] but in contrast to a report implicating DCs in negative selection of self-reactive thymocytes [29]. If CD11c^+^ DCs were required in negative selection of PLP-reactive thymocytes, we would have expected significantly increased frequencies of TCR transgenic CD4^+^ T cells in DC deficient PLP TCR Tg. CD11c-DTA SJL than in PLP TCR Tg. DTA control mice, which we did not observe. Notably, CD4^+^T cells that had developed in the absence of DCs in PLP TCR Tg. CD11c-DTA SJL were not impaired in their pathogenic function because they mediated severe EAE in RAG2 deficient SJL recipients as efficiently as those from PLP TCR Tg. DTA SJL mice.

In the absence of DCs, spontaneous EAE developed significantly earlier in PLP TCR Tg. CD11c-DTA SJL mice as compared to PLP TCR Tg. DTA control mice. Despite the lack of DCs, activation of naïve PLP-specific CD4^+^ T cells in PLP TCR Tg. CD11c-DTA SJL mice was not impaired but instead resulted in increased frequencies of effector memory CD4^+^ T cells than in DC sufficient control mice. These results are in contrast to findings in a spontaneous murine lupus model that showed constitutive deletion of DCs ameliorated disease and did not affect activation of T cells [42]. The reason for the different findings between the two studies are unclear but may be due to differences between the pathogenesis of lupus and EAE, as well as genetic differences between the lupus-prone MRL-MpJ-*Fas^lpr^*/J strain and the EAE prone SJL/J strain. Our findings indicate that non-DCs were capable to compensate for the loss of DCs by efficiently presenting endogenous autoantigen and priming encephalitogenic T cells in PLP TCR Tg. CD11c-DTA SJL mice. A number of mechanisms may explain the increased frequency of activated CD4^+^ T cells and accelerated EAE onset in PLP TCR Tg. CD11c-DTA SJL mice, including the loss of tolerogenic DCs [10]. Thus, depletion of tolerogenic DCs may decrease tolerance to self-antigens, including PLP, resulting in enhanced priming of PLP-specific T cells and accelerated EAE in PLP TCR Tg. CD11c-DTA SJL mice.

A number of laboratories have investigated the role of DCs in models of actively or passively induced EAE. A pioneering study by Dittel et al. reported that DCs pulsed with encephalitogenic peptide Ac_1–11_ derived from myelin basic protein (MBP), were sufficient to initiate EAE when co-transferred with CD4^+^ T cells from Ac_1–11_ TCR transgenic mice [18]. We previously demonstrated that activation of DCs by microbial TLR ligands was sufficient to break CD4^+^ T cell tolerance and trigger EAE in PLP TCR transgenic mice in an EAE resistant strain [17]. Greter and colleagues reported that Flt-3 ligand expanded DCs were able to present autoantigen and prime myelin-reactive T cells in vivo, resulting in EAE exacerbation [19]. Furthermore, studies in Steve Miller’s laboratory showed that DCs in the inflamed CNS activated myelin-specific Th cells specific for different PLP epitopes during the course of EAE and drove Th17 cell differentiation [33] [20]. Collectively, these studies demonstrated the ability of DCs to promote myelin-reactive T cell responses resulting in active or passive EAE. Conversely, other studies reported that DCs in the periphery or within the CNS were able to protect mice from EAE induction [21–23]. The reason for these opposite roles of DCs in EAE pathogenesis is unclear but it may at least partially be associated with differences in the subsets and/or activation state of the examined DCs.

Since these studies, a number of genetically engineered mouse models that allow for conditional selective ablation of DCs, including CD11c-DTR mice, have become available and have been utilized to further interrogate the role of DCs in immune responses [43]. To elucidate the requirement of DCs in active and passive EAE following conditional depletion of CD11c^+^ DCs, we generated novel CD11c-DTR SJL and CD11c-DTR SJL BM chimeric mice, respectively. To our knowledge, these novel mice allowed selective ablation of CD11c^+^ DCs in autoimmune-prone SJL mice for the first time. Short term and long term administration of diphtheria toxin (DT) extensively depleted cDCs in CD11c-DTR SJL and CD11c-DTR SJL BM chimeric SJL mice, respectively, which was consistent with findings from studies in the non-autoimmune prone C57BL/6 mice [12]. Notably, DT administration also partially depleted pDCs in CD11c-DTR SJL mice, which was not observed in CD11c-DTR C57BL/6 mice [24, 44]. It is conceivable that higher level of CD11c expression in pDCs of SJL mice than in C57BL/6 mice may have contributed to increased efficiency of DT induced ablation of pDCs in CD11c-DTR SJL mice.

Because of their potent function as APCs, it is generally assumed that CD11c^+^ DCs are required for efficient priming and differentiation of myelin-specific CD4^+^ T cells following myelin antigen immunization. However, we show that PLP TCR transgenic CD4^+^ T cells transferred to CD11c-DTR SJL mice or DC sufficient control mice proliferated comparably in draining lymph nodes in response to cognate antigen immunization. Furthermore, our study shows that DCs were dispensable for differentiation of PLP TCR-transgenic CD4^+^ T cells to Th1 or Th17 cells, both of which play decisive roles in EAE pathogenesis [45, 46]. These findings support a non-essential role of DCs in the priming of self-reactive CD4^+^ T cells, which agrees with the results reported in DC depleted mice in MOG-induced EAE or in murine lupus [24, 25, 42].

Transfer of purified CD4^+^ T cells from naïve PLP TCR transgenic SJL mice into lymphopenic syngeneic recipient mice results in rapid development of EAE [32]. Here, we show that long-term DT induced depletion of DCs in CD11c-DTR *Rag2*−/− SJL BM chimeras significantly accelerated EAE onset mediated by adoptively transferred PLP TCR transgenic CD4^+^ T cells. This finding indicates that in the absence of DCs, other professional APCs such as B cells or macrophages were able to present CNS myelin antigen to myelin-specific T cells. Since CD11c-DTR *Rag2*−/− BM chimeras genetically lack B cells [47], macrophages or other CD11c^−^ myeloid cells may have compensated for the loss of DCs in these mice and promoted encephalitogenic T cell responses.

Although constitutive or conditional ablation of DCs was consistently profound in our models, we cannot exclude the possibility that low numbers of remaining DCs may have been sufficient to promote the potent PLP-specific T cell responses observed in DC depleted mice. However, we think this is unlikely because the onset of both spontaneous EAE and passive EAE was significantly accelerated in DC ablated mice as compared to DC sufficient control mice. If DCs had been essential for the priming of PLP-specific CD4^+^ T cells, we would have expected to observe impaired activation or priming of pathogenic CD4^+^ T cells resulting in delayed onset of EAE in DC ablated mice. Instead, T cell activation and EAE onset were significantly enhanced in DC depleted mice, suggesting that DCs contributed to maintenance of self-tolerance to PLP autoantigen.

These findings are consistent with a study providing evidence that DCs played a protective role in EAE induced by MOG immunization or transfer of MOG-primed T cells [25]. The mechanism by which DCs apparently exert tolerogenic effects in PLP-induced EAE in SJL mice remains to be determined. Potential mechanisms may include DC-mediated induction of regulatory T cells, which was implied in one [25] but not in another EAE study [24] in CD11c-DTR C57BL/6 mice.

In conclusion, utilizing our newly established constitutive or conditional DC ablation models in autoimmune prone SJL mice, we demonstrate that development, selection and activation of PLP-specific CD4^+^ T cells was not impaired in the absence of CD11c^+^ DCs. Instead, non-DCs compensated for the loss of DCs by efficiently promoting pathogenic Th responses, resulting in significantly accelerated onset of spontaneous or passive EAE in DC depleted mice. Our results indicate that CD11c^+^ DCs regulate PLP-specific T cell responses in EAE, which validates DCs as mediators of T cell tolerance to CNS myelin antigens in SJL mice.

## Acknowledgments

We thank Aaron Wexler for help in initial backcrossing of mice used in this study. This work was supported by Pennsylvania State University College of Medicine and the H. G. Barsumian Fund to HW.

